# Leptin receptor expression in the dorsomedial hypothalamus stimulates breathing during NREM sleep in *db/db* mice

**DOI:** 10.1101/2020.09.29.318758

**Authors:** Huy Pho, Slava Berger, Carla Freire, Lenise J Kim, Mi-Kyung Shin, Stone R Streeter, Nishitha Hosamane, Meaghan E Cabassa, Frederick Anokye-Danso, Olga Dergacheva, Mateus Amorim, Thomaz Fleury-Curado, Jonathan J Jun, Alan R Schwartz, Rexford S Ahima, David Mendelowitz, Vsevolod Y Polotsky

## Abstract

Obesity can lead to recurrent upper airway obstruction (obstructive sleep apnea, OSA) during sleep as well as alveolar hypoventilation. We have previously shown that leptin stimulates breathing and treats OSA in leptin-deficient *ob/ob* mice and leptin-resistant diet-induced obese mice. Our previous data also suggest that leptin’s respiratory effects may occur in the dorsomedial hypothalamus (DMH). We selectively expressed leptin receptor *LepR^b^* in the DMH neurons of obese *LepR^b^*-deficient *db/db* mice (*LepR^b^*-DMH mice), which hypoventilate at baseline, and showed that intracerebroventricular injection of leptin in these animals increased inspiratory flow, tidal volume and minute ventilation during NREM sleep without any effect on the quality of NREM sleep or CO_2_ production. Leptin had no effect on upper airway obstruction in *LepR^b^-DMH* animals. We conclude that leptin stimulates breathing and treats obesity related hypoventilation acting on LepR^b^-positive neurons in the DMH.

## INTRODUCTION

Obesity is a highly prevalent condition observed in 34.9% of US adults^1^. Obesity causes sleep disordered breathing (SDB), which can be manifested by obstructive sleep apnea (OSA) and obesity hypoventilation syndrome (OHS). OSA is the most prevalent SDB, affecting 50% of obese patients^2–5^. OSA is characterized as recurrent obstructions of the upper airway during sleep, leading to intermittent hypoxia, sleep fragmentation and intrathoracic pressure swings^6,7^. OHS is defined by sleep-related decreases in ventilatory drive, leading to daytime hypercapnia and hypoventilation during sleep in obese individuals and it has been reported in 10-20% of obese patients with OSA^8^. Continuous positive airway pressure (CPAP) is an efficacious treatment for both OSA^9,10^ and OHS^11^. Nevertheless, poor adherence to CPAP^12^ limits its therapeutic use and emphasizes the unmet need for pharmacotherapy development. Pharmacological development in SDB has been hindered by the lack of rodent models of OSA.

We developed and validated novel plethysmographic methods for monitoring high-fidelity airflow and respiratory effort signals *continuously* during sleep in mice^13^. Upper airway obstruction was defined by the presence of inspiratory airflow limitation characterized by an early inspiratory plateau in airflow at a maximum level (V_I_max) while effort continued to increase^13–16^. Moreover, we have demonstrated that obesity plays a major role in the pathogenesis of inspiratory flow limitation and OSA in mice and human alike^17–23^.

Leptin is an adipocyte-produced hormone that regulates food intake and metabolic rate^24–26^. Leptin also plays an important role in the control of breathing^27^ and upper airway patency^20,28–30^ during sleep^18,19^. Leptin-deficient *ob/ob* mice develop SDB, characterized by **(a)** hypoventilation during sleep and the elevated partial pressure of carbon dioxide in the arterial blood (PaCO_2_) ^27^; **(b)** inspiratory flow limitation and recurrent hypopneas, similar to human OSA, which were reversed by leptin infusion^18,19^. However, hyperleptinemia is a common feature of human obesity^29,30^ and obese humans are resistant to metabolic effects of leptin. Moreover, OSA and obesity hypoventilation are also associated with leptin resistance^28,31,32^ and non-invasive ventilation decreases both CO_2_ levels and leptin levels, independent of obesity^33^. The blood-brain barrier is one of the key sites of leptin resistance^34,35^. Leptin delivery beyond the blood brain barrier effectively reverses OSA and hypoventilation in both leptin-deficient and leptin-resistant mice^19,20,36^. Leptin signals *via* the long isoform of leptin receptor, LepR^b^^37–40^, which is ubiquitous in the hypothalamus and many areas of medulla^41^. However, localization of respiratory effects of leptin and identity of leptin-responsive respiratory neurons remain unknown.

We have previously injected leptin into the lateral and fourth cerebral ventricles of *ob/ob* mice and found that leptin’s effects on upper airway patency likely occur in the forebrain rather than in the medulla. We also found that hypoglossal motoneurons are synaptically connected to the dorsomedial hypothalamus (DMH), which abundantly expresses LepR^b^^19^. Notably, the DMH abundantly expresses another important metabolic receptor, melanocortin 4 receptor (MC4), which may mediate leptin’s effect^42^. MC4 deficiency leads to severe obesity^43,44^ and has been associated with SDB^45^. We hypothesized that leptin’s effect on SDB may occur via LepR^b^ signaling in the DMH. In order to examine this hypothesis we expressed *LepR^b^* exclusively in the DMH of obese *LepR^b^-* deficient *db/db* mice and performed sleep studies after transgene expression at baseline and after intracerebroventricular (ICV) leptin infusion.

## METHODS

### Animals

In total, 46 male homozygous B6.BKS(D)-*Lep^db^*/J (*db/db*) leptin receptor deficient mice from Jackson Laboratory (Bar Harbor, ME, Stock #000697) were used for this study. *LepR^b^-GFP* mice (n=4) generated by breeding *LepR^b^-Cre* [B6.129(Cg)-*Leprtm2(cre)*Rck/J, Stock #008320] and green fluorescent protein (GFP) floxed mice [B6.129-Gt(ROSA)26Sortm2Sho/J, Stock #004077] from the Jackson Laboratory were used for histology only. Water and food were available *ad libitum*. Mice were housed at a 12h-light/dark cycle (7am–7pm lights on), and temperature of 26°C. Food consumption and body weight were monitored daily throughout the sleep study protocol. All protocols were approved by the Johns Hopkins University Animal Care and Use Committee (ACUC) and all animal experiments were conducted in accordance with ACUC guidelines.

### Arterial Blood Gas

Seven *db/db* mice had an arterial catheter implanted in the left femoral artery under 1-2% anesthesia as previously described^14,46^. In brief, a small incision was made to expose the femoral artery and an arterial catheter was placed 5-8 mm deep into the femoral artery. The catheter was glued in place and fed subcutaneously to be dorsally attached to a single channel fluid swivel (model 375/25, Instatech Laboratories, Plymouth Meeting, PA, USA), which slowly perfused a heparin saline solution (1000 U heparin/L saline) *via* an infusion pump (0.5 mL/day). Mice recovered for 48-72 hours. Arterial blood gas was analyzed in awake unrestrained unanesthetized mice by removing 150 μL of arterial blood, which was tested in a blood gas analyzer (Radiometer ABL 800 Flex, Diamond Diagnostics, Holliston, MA, USA).

### Sleep Studies

A two-arm crossover study design was used **(Figure 1).** Mice were randomized to receive stereotactic injections of *Ad-LepR^b^* or *Ad-mCherry*. After the 9-day gene expression period, each mouse was recorded with treatment of either ICV vehicle (phosphate buffered saline, PBS) or ICV leptin. Polysomnography was performed in 26 mice, aged 15 weeks, at two time points with a 72-hour washout period between studies.

**Figure 1.**
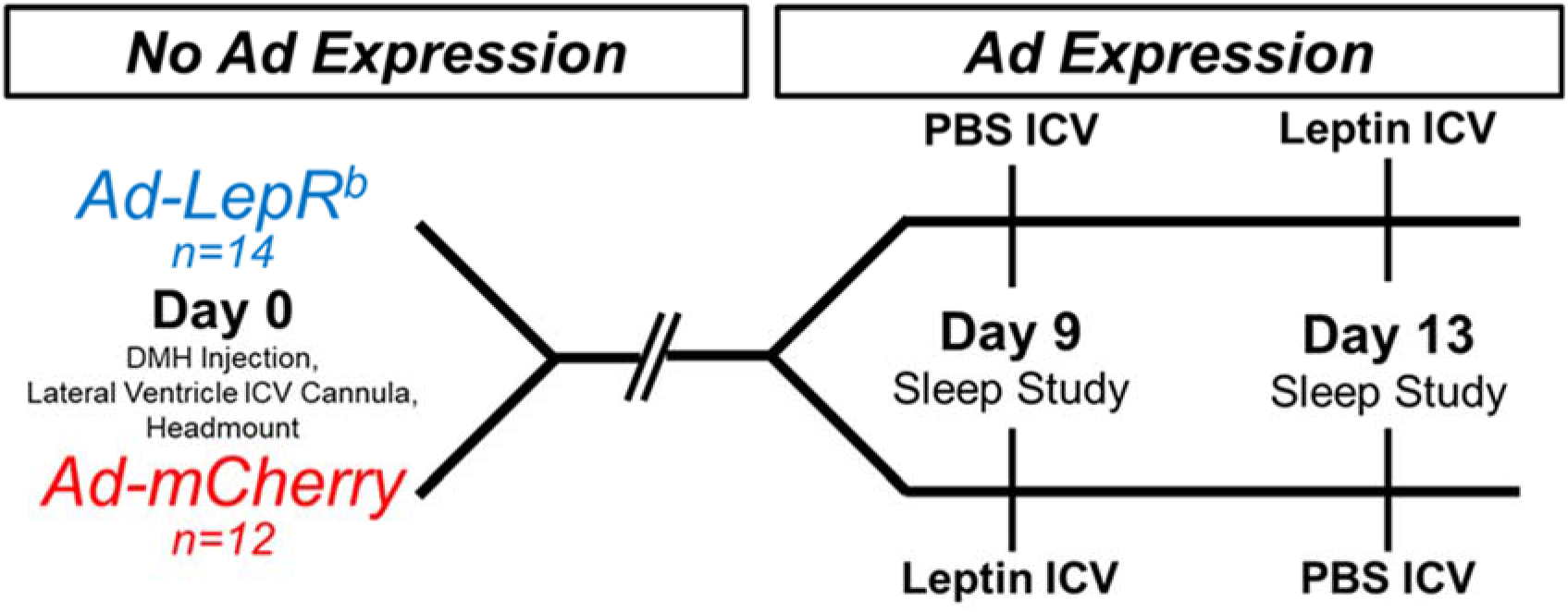
Experimental study design. LepR^b^-deficient db/db mice were randomly transfected with Ad-LepR^b^ or Ad-mCherry in the dorsomedial hypothalamus (DMH). Mice received intracerebroventricular (ICV) infusions of either PBS or leptin in a cross-over design and underwent polysomnography at days 9 and 13 after injection. LepR^b^: long isoform of leptin receptor.

Mice were anesthetized with isoflurane 1-2% and placed in the stereotaxic system (Model 963 with 923-B Head Holder, David Kopf Instruments, Tunjunga, CA) and ~125μL bilateral viral injections of either *Ad-LepR^b^* (ADV-263380, Vector Biosystems, 2-5×10^10^ PFU/mL) or *Ad-mCherry* (Cat No: 1767, Vector Biolabs, 1×10^10^ PFU/mL) were performed using −1.88 mm caudal, ±0.40 mm lateral, −5.00 mm ventral as DMH coordinates from the bregma. To allow ICV administration of leptin or vehicle, a cannula (PlasticsOne, Roanoke, VA) was implanted in the right lateral ventricle (LV) at −0.60 mm caudal, −1.20 mm lateral, −3.00 mm ventral from the bregma. Headmount procedure was performed immediately following viral transfection and ICV cannula placement. Custom 6-pin headmounts (8231-SM-C, Pinnacle Technology, Lawrence, KS) were implanted for electroencephalogram (EEG) and electromyogram (EMG) recordings. Briefly, three holes were bored through the skull in the left and right frontal regions and left parietal region to allow implantation of 0.10” silver electrodes with wire leads (no. 8403, Pinnacle Technology). The EEG leads from the custom headmount were twisted together with the three silver electrodes and coated with silver conductive epoxy (no. 8331, MG Chemicals) to provide unipolar conductive EEG electrodes. EMG leads were tunneled subcutaneously and placed over the nuchal muscle posterior to the skull. Dental acrylic (Lang Dental, Wheeling, IL) was used to secure the headmount and cannula in place.

For polysomnography we used a modified whole body plethysmography (WBP) chamber system to measure tidal airflow and sleep-wake state continuously, generating high-fidelity tidal volume and airflow signals, as previously described^13^. In brief, following ICV treatment of PBS (pH 7.4, 2 μL) or leptin (10 μg in 2 μL of PBS), mice were placed in the WBP chamber to be recorded from 10:00am to 4:00pm. Mice were weighted prior to sleep recording. Mouse rectal temperature was measured and averaged between beginning and end of sleep study. Mice were acclimated to the chamber prior to recording for at least 3 days, 1h per day. During full polysomnographic recordings, the chamber was humidified to ~90% relative humidity and ~29°C while a slow leak allowed atmospheric pressure equilibrium. The WBP’s reference chamber filtered out ambient noise from the pressure signal acquired by a transducer (Emka Technologies). Positive and negative pressure sources were utilized in series with mass flow controllers (Alicat Scientific) and high-resistance elements to generate a continuous bias airflow through the animal chamber while maintaining a sufficiently high time constant. Tidal airflow was calculated from the plethysmography chamber pressure signal using the Drorbaugh and Fenn equation^47^, which required the measurements of mouse rectal temperature, chamber temperature, room temperature, relative humidity, and chamber gas constant, calculated by utilizing a known volume injection and the resultant chamber pressure deflection. The tidal volume signal was differentiated electronically to generate an airflow signal.

All signals were digitized at 1,000 Hz (sampling frequency per channel) and recorded in LabChart 7 Pro (Version 7.2, ADInstruments, Dunedin, NZ). Sleep-wake state was scored visually in 5 second epochs based off standard criteria of EEG and EMG frequency content and amplitude, as previously described^14,18,19^. Wakefulness was characterized by low-amplitude, high-frequency (~10 to 20 Hz) EEG waves and high levels of EMG activity compared with the sleep states. NREM sleep was characterized by high-amplitude, low frequency (~2 to 5 Hz) EEG waves with EMG activity considerably less than during wakefulness. REM sleep was characterized by low-amplitude, mixed frequency (~5 to 10 Hz) EEG waves with EMG amplitude either below or equal to that during NREM sleep. Respiratory signals were analyzed from all REM sleep periods and from periods of NREM sleep sampled periodically at 20-second stretches every half an hour throughout the total recording time. Custom software was used to demarcate the start and end of inspiration and expiration for subsequent calculations of timing and amplitude parameters for each respiratory cycle.

We utilized each breath’s respiratory characteristic to describe maximal inspiratory airflow (V_I_max) and components of minute ventilation (V_E_). We developed an algorithm using the airflow and respiratory effort signals to determine if a breath was classified as inspiratory airflow limited, defined by an early inspiratory plateau in airflow while effort continued to increase. The software provided peak flow values during the first half (V_I_max1), midpoint (V_I_50), and second half (V_I_max2) of inspiration. Breaths resembling sniffs were initially defined as nonflow limited by their short duration, having an inspiration time with a z-score lower than 1.75. Breaths having sufficient inspiration time were then classified as inspiratory flow limited if a mid-inspiratory flow plateau was present^14,18,19^.

### Metabolic measurements

Metabolic studies were performed in a separate subset of mice (n=4 in the *Ad-LepR^b^* group and n=5 in the *Ad-mCherry* group) according to the design described for the sleep studies starting nine days after viral injection and cannula implantation **(Figure 1)**, except that the headmount was not installed and the washout period was 1 week between studies. Mice were placed in individual Comprehensive Laboratory Animal Monitoring System (CLAMS) units (Oxymax series; Columbus Instruments, Columbus, OH) for a 24-hour acclimation period followed by 48 hours of continuous recordings. The 48-hour recordings started at 10:00 am with 4 brief interruptions (daily at 10:00 am and 6:00 pm) for ICV injections. Each treatment infusion was 2 μL of PBS or leptin (10μg/2μL, R&D Systems, Minneapolis, MN). Data collected 30 minutes prior and 30 minutes after the ICV injections were excluded from the analysis. The CLAMS units were sealed and equipped with O_2_ electrochemical sensors, CO_2_ infrared sensors and infrared beam movement sensors. Consumed O_2_ (VO_2_) and produced CO_2_ (VCO_2_) were collected every 11 minutes and measurements were utilized to calculate the respiratory exchange ratio (RER). Motor activity was quantified by the number of infrared beam interruptions. Total horizontal and vertical beam breaks were summed and presented as motor activity. Metabolic cages were kept in a 12 h light / dark cycle (7am-7pm lights on) with food and water *ad libitum* and a consistent environmental temperature of 24°C.

### Immunofluorescence

In *db/db* mice (n=4) and *LepR^b^-GFP* mice (n=4) histology was performed upon completion of physiology experiments. Mice were anesthetized with isoflurane 1-2% and rapidly perfused with PBS followed by ice-cold 4% paraformaldehyde in distilled water. The brains were carefully removed, postfixed in 4% paraformaldehyde for 1h at 4°C, and cryoprotected in 20% sucrose in PBS overnight at 4°C. Then, brains were covered with O.C.T. Compound (Tissue Tek, Cat No: 4583) and frozen using 2-methylbutane on dry ice. Frozen brains were cut into 16 μm thick coronal sections on a sliding microtome and stored at −20°C until further use. Immunohistochemical staining of brain slices was performed as described previously^20,46^ with modifications. Briefly, sections were blocked for 2h with 10% normal goat serum in PBS/0.5% TritonX-100 (PBST). The antibodies were added and incubated overnight at 4°C. The next day, sections were washed with PBST at room temperature and incubated for 2h with secondary antibodies. The sections were washed again with PBST and cover-slipped with mounting medium for fluorescence with DAPI (4,6-Diamidino-2-phenylindole, Vectashield, California, USA).

LepR^b^ protein expression in the hypothalamus of *db/db* mice after infection with *Ad-LepR^b^-GFP* and *LepR^b^-GFP* mice was detected based on the presence of a reporter, enhanced green fluorescent protein (EGFP)^41^. Positive expression of leptin receptor in the *Ad-LepR^b^*-treated *db/db* mice was confirmed by immunostaining with chicken anti-rat *LepR^b^* antibody (1:100, CH14104, Neuromics, MN) and detected with goat anti-Chicken IgY secondary antibody, Alexa Fluor 568 (1:500, Cat No: A-11041).

Phenotypic identification of the LepR^b^-positive cells in the hypothalamus of the *Ad-LepR^b^-GFP* treated *db/db* mice and *LepR^b^-GFP* mice was assessed by immunostaining for Anti-NeuN as a neuronal marker (1:100, ab104225, ABCAM, Cambridge, MA), anti-Glial Fibrillary Acidic Protein (GFAP) as an astrocyte marker, (1:100, N1506, DAKO, Carpinteria, CA), and anti-ionized calcium binding adaptor molecule 1 (Iba1) as a microglia marker, (1:100. ab178846, ABCAM, Cambridge, MA). Then the slides were incubated with goat anti-rabbit secondary antibody, Alex Fluor 647 (1:500, Cat No: A-21245). Co-localization with MC4 was examined with rabbit polyclonal to MC4 antibodies (1:100, ab24233, ABCAM, Cambridge, MA).

Fluorescence images were examined with an Inverted Axio Observer 3 microscope (Carl Zeiss, Jena, Germany) equipped with an Axiocam 512 camera. The following filters have been used: for EGFP the led module-475 nm filter; for Alexa 647 the led module-630 nm filter, for Alexa 568 the led module-567 nm filter, and for DAPI, the led module-385 nm filter. All the images were processed and merged with the softwares Carl Zeiss Image and ImageJ (NIH). At least 300 cells per mouse were manually counted and the percentage of LepR^b^-positive cells was calculated.

### Statistical Analysis

Data were tested for normality using Shapiro-Wilk’s test. Effects of virus transfection (*Ad-LepR^b^* vs *Ad-mCherry*) and treatment (leptin vs PBS), as well as their interaction (virus*treatment), on sleep studies and metabolic parameters were verified using Generalized Estimating Equations (GEE). GEE was used to obtain adjusted estimates of association between repeated measurements within subjects and also between the groups at each time point^48^. Pairwise comparisons were performed by Bonferroni’s post hoc test. The effects of treatment randomization according to the crossover design were firstly tested for each dependent variable and no statistical significance was observed. The goodness of fit of each model was assessed by Quasi-likelihood under Independence Model Criterion (QIC) and the residuals were tested for normality using Q-Q plots. Statistical analyses were performed in SPSS version 20.0 (IBM SPSS Inc, Chicago, IL) and the data are represented as mean±SEM. Statistical significance was considered at a level of p<0.05.

## RESULTS

### Arterial blood gas analysis

Arterial blood gas in *db/db* mice at baseline was pH 7.37 ± 0.01; PaCO_2_ 42.3 ± 2.1 mmHg, and PaO_2_ 79.8 ± 6.1 mmHg. PaCO_2_ was markedly higher and pH was lower than we previously reported in lean C57BL/6J mice (pH 7.46 ± 0.06, PaCO_2_ 30 ± 5.0 mmHg, and PaO_2_ 98 ± 3.9 mmHg)^14^.

### Leptin receptor expression in the DMH

LepR^b^ and mCherry were successfully expressed in the DMH of *db/db* mice transfected with *Ad-LepR^b^* (**Figure 2A-C**) and *Ad-mCherry* (**Supplemental Figure 1**). Expression of LepR^b^ or control mCherry was not detected in any other area of the brain. Expression of LepR^b^ in the DMH was confirmed both by LepR^b^ antibody staining (**Figure 2A, B**) and by GFP fluorescence (**Figure 2C**). Quantitative analysis of GFP fluorescence in *db/db* mice showed that *Ad-LepR^b^* was present in 11.6 ± 1.7 % of all DMH cells. LepR^b^ were identified in neurons by co-localizing *Ad-LepR^b^-GFP* and NeuN (**Figure 2C**). LepR^b^ was absent in astrocytes and microglia as shown by the lack of co-localization of *Ad-LepR^b^-GFP* with GFAP (**Figure 2D**) and IBA-1 (**Figure 2E**), respectively. Nearly all *Ad-LepR^b^* positive cells were MC4-positive as it was evident from co-localization of MC4 red and GFP resulting in orange color (**Figure 2F**). Quantitative analysis of GFP fluorescence in DMH of *LepR^b^-Cre-GFP* mice showed that LepR^b^ was present in a similar pattern as in *Ad-LepR^b^-GFP*-transfected *db/db* mice: LepR^b^ was expressed in a similar percentage of cells, 9.6 ± 0.8 %, and was observed exclusively in neurons (**Figure 2G**), but not in astrocytes (**Figure 2H**) or microglia (not shown). Co-localization of MC4 and LepR^b^-GFP was detected in some cells shown with arrows. Higher magnification (**Figure 2I**, insert) showed that LepR^b^ and MC4 colocalized in a majority of cells, but LepR^b^ was distributed diffusely throughout the cell, whereas MC4 was detected perinuclearly (merged blue DAPI + red MC4 resulting in purple color) suggesting that it was inactive^49^. Thus, we successfully expressed LepR^b^ in the DMH and the distribution of LepR^b^ in MC4 positive neurons resembled the natural distribution of LepR^b^ in *LepR^b^-Cre-GFP* mice.

**Figure 2.**
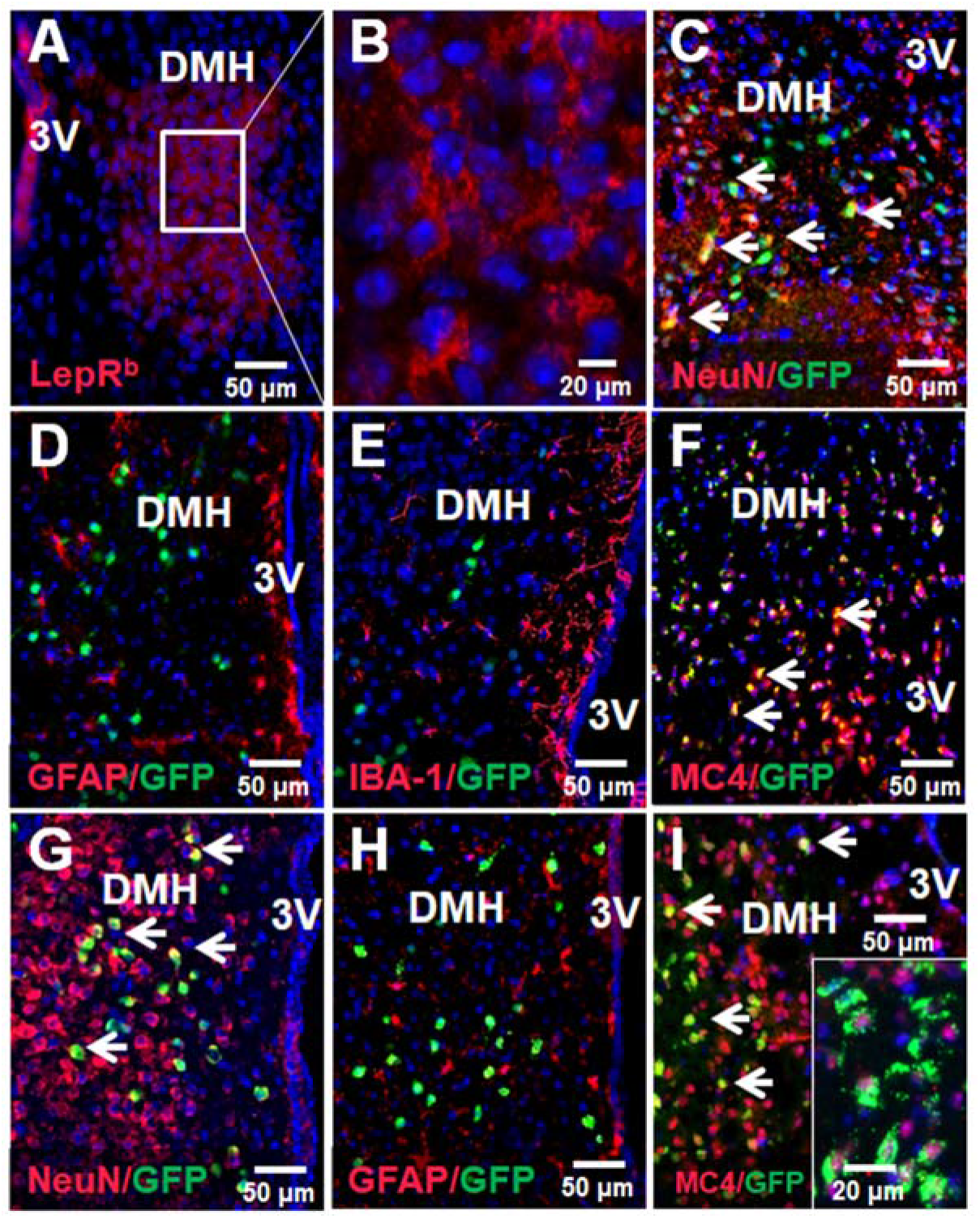
Localization of LepR^b^ in the DMH of db/db mice after Ad-LepR^b^ infection (A-F) compared to LepR^b^ in DMH of LepR^b^-Cre-GFP mice (G-I). (A) LepR^b^ (in RED) was expressed in the DMH of db/db mice 9 days after Ad-LepR^b^ infection; DAPI (4’,6-diamidino-2-phenylindole) in BLUE. (B) same at higher power; (C) GFP in DMH of db/db mice was localized in neurons (NeuN, RED) as evidenced by ORANGE color after merging (arrows), but not in (D) astrocytes (glial fibrillary acidic protein, GFAP, RED) (E) or microglia (IBA-1101, RED); (F) in db/db mice GFP co-localized with melanocortin 4 receptor (MC4). In LepR^b^-Cre mice LepR^b^ localized in neurons (G, ORANGE), but not in astrocytes (H). (I) LepR^b^ colocalized with MC4 (ORANGE at lower power and co-localization at higher power, corner insert); note perinuclear location of MC4 (insert), which is typical in the inactive state^49^. 3V, third ventricle.

### Basic characteristics of db/db mice

Characteristics of *db/db* mouse that underwent sleep recordings are shown in **Table 1**. Weight, age, body temperature, and food intake were not significantly different between *Ad-LepR^b^* and the *Ad-mCherry* groups at the time of transfection.

**Table 1.** Characteristics of age, weight, temperature, and food intake for db/db mice transfected with either Ad-mCherry or Ad-LepR^b^ and infused with PBS or Leptin ICV. Weight was recorded prior to sleep recording. Rectal temperature was averaged between beginning and end of sleep recording. Food intake was measured for 24h post leptin or PBS treatment. Values presented as mean values ± standard error and range. *p<0.05 ***p<0.001

|  | n | Age, weeks |  | Weight, g |  | Temperature, °C |  | Food Intake, g |  |
| --- | --- | --- | --- | --- | --- | --- | --- | --- | --- |
|  |  | Mean ± SE | Range | Mean ± SE | Range | Mean ± SE | Range | Mean ± SE | Range |
| ***Ad-mCherry*** |  |  |  |  |  |  |  |  |  |
| PBS | 12 | 15.9 ± 0.0 | 15.0-17.0 | 47.4 ± 0.9 | 39.4-50.6 | 36.1 ± 0.3 | 34.6-37.5 | 3.0 ± 0.5 | 2.1-5.9 |
| Leptin | 12 | 16.0 ± 0.1 | 15.0-17.1 | 46.9 ± 1.0 | 39.9-51.5 | 35.4 ± 0.2 | 34.4-36.6 | 3.0 ± 0.4 | 2.3-5.2 |
| ***Ad-LepR<sup>b</sup>*** |  |  |  |  |  |  |  |  |  |
| PBS | 14 | 15.7 ± 0.1 | 14.0-17.4 | 49.4 ± 1.3 | 35.7-55.4 | 35.8 ± 0.3 | 33.3-37.8 | 3.2 ± 0.2 | 1.6-4.5 |
| Leptin | 14 | 15.7 ± 0.1 | 14.0-17.0 | 49.8 ± 1.2 | 36.9-56.0 | 36.3 ± 0.2* | 35.1-38.2 | 2.5 ± 0.3*** | 1.6-3.8 |

Body temperature increased (p<0.05) and food intake decreased (p<0.001) in the *Ad-LepR^b^* group when treated with leptin compared to PBS.

### Effect of leptin signaling in DMH on sleep architecture

Sleep architecture for all groups is described in **Table 2**. There was no significant difference in sleep efficiency, total sleep time, NREM sleep time, NREM sleep bout number or bout length between *Ad-LepR^b^* and *Ad-mCherry* groups at baseline, nor was there an effect of ICV leptin. Duration of REM sleep was lower in *Ad-LepR^b^* transfected mice treated with leptin compared to PBS (p<0.001) and compared to *Ad-mCherry* treated with leptin (p<0.001). There was no significant difference in REM sleep bout length between any of the groups, but REM sleep bout number was significantly lower after leptin treatment in mice transfected with *Ad-LepR^b^* (p<0.001).

**Table 2.** Sleep characteristics for db/db mice transfected with either Ad-mCherry or Ad-LepR^b^ and infused with PBS or Leptin ICV. TST: total sleep time. Values presented as mean values ± standard error. *p<0.05 **p<0.01 ***p<0.001

|  | n | Sleep Efficiency | Sleep Time (min) |  |  | Sleep Bout Number (n) |  | Sleep Bout Length (s) |  |
| --- | --- | --- | --- | --- | --- | --- | --- | --- | --- |
|  |  | (%) | TST | NREM | REM | NREM | REM | NREM | REM |
| *Ad-mCherry* |  |  |  |  |  |  |  |  |  |
| PBS | 12 | 73.6 ± 4.3 | 273.0 ± 17.6 | 268.6 ± 17.2 | 4.5 ± 2.2 | 41.9 ± 4.9 | 4.7 ± 2.1 | 486.2 ± 96.9 | 44.2 ± 10.5 |
| Leptin | 12 | 72.3 ± 4.0 | 281.2 ± 20.5 | 274.8 ± 21.1 | 6.5 ± 1.4 | 31.4 ± 6.0 | 5.5 ± 1.0 | 446.8 ± 91.9 | 72.5 ± 10.4 |
| *Ad-LepR<sup>b</sup>* |  |  |  |  |  |  |  |  |  |
| PBS | 14 | 64.3 ± 3.7 | 241.0 ± 16.6 | 235.9 ± 16.5 | 5.2 ± 0.8 | 57.8 ± 5.3 | 5.0 ± 0.7 | 301.0 ± 50.9 | 65.9 ± 4.9 |
| Leptin | 14 | 69.6 ± 3.9 | 267.1 ± 19.1 | 266.1 ± 19.2 | 1.0 ± 0.3*** | 58.5 ± 8.3 | 1.3 ± 0.4*** | 447.6 ± 115.6 | 34.2 ± 9.6 |

Representative polysomnography recordings of NREM sleep in *db/db* mice transfected with either *Ad-LepR^b^* or *Ad-mCherry* during PBS and leptin treatments are shown in **Figure 3**. Leptin treatment did not change the respiratory pattern in *Ad-mCherry* transfected mice (**Figure 3**, Left Panels). In contrast, ventilation visually increased after leptin administration in the *Ad-LepR^b^* transfected mice (**Figure 3**, Right Panels). The respiratory pattern in REM sleep was not affected by types of virus or injection (**Supplemental Figure 2**).

**Figure 3.**
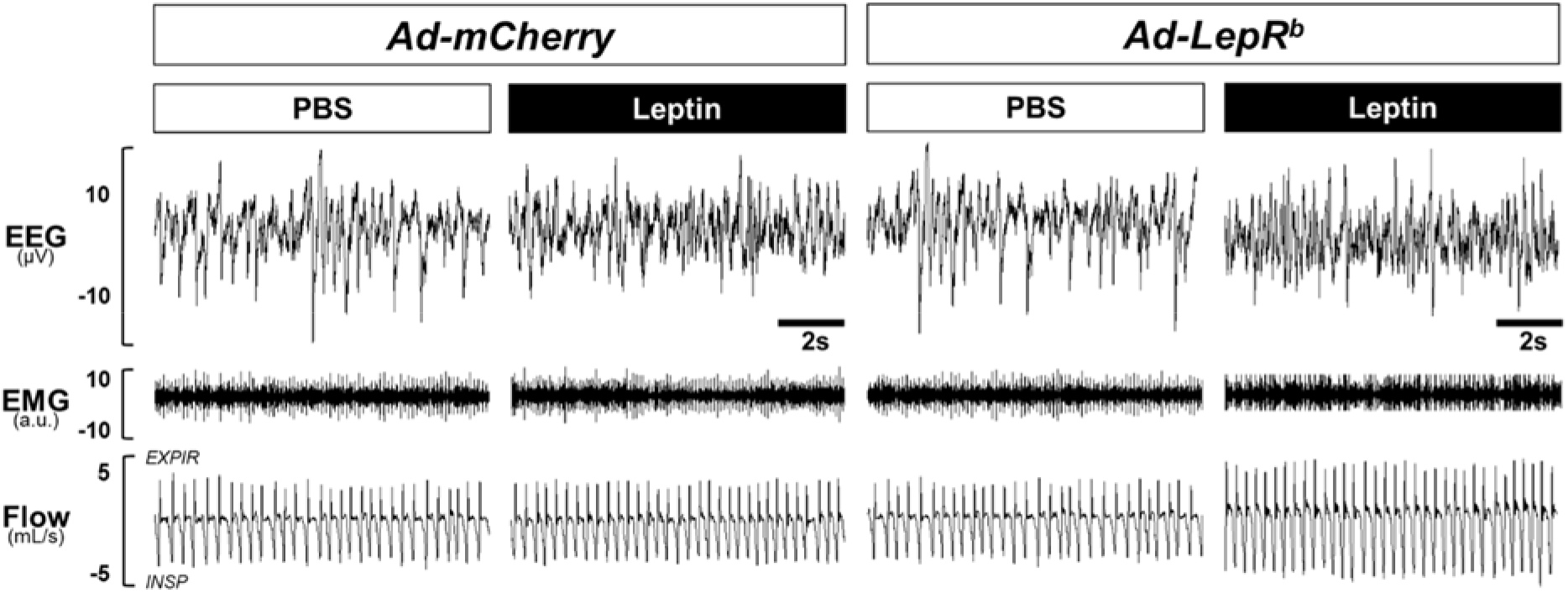
Representative NREM sleep recordings in db/db mice transfected with (A) Ad-mCherry or (B) Ad-LepR^b^ in the dorsomedial hypothalamus. Intracerebroventricular administration of leptin augmented ventilation in Ad-LepR^b^ mice without altering sleep architecture. No leptin effect was observed after Ad-mCherry transfection.

### Effect of leptin on non-flow limited breathing during sleep

In mice transfected with *Ad-LepR^b^*, administration of leptin to the lateral ventricle increased minute ventilation (nV_E_) during NREM sleep from 0.78 ± 0.06 to 0.97 ± 0.07 mL/min/g, p<0.01, whereas no effect was observed in mice transfected with *Ad-mCherry* (0.72 ± 0.06 mL/min/g with PBS and 0.70 ± 0.07 mL/min/g with leptin) (**Figure 4A**). This 24% increase in minute ventilation was attributed to an increase in tidal volume (V_T_) from 0.19 ± 0.01 mL to 0.22 ± 0.02 mL (p<0.001), whereas respiratory rate (RR) was unchanged (**Figure 4B-C**). Leptin did not affect minute ventilation in REM sleep in either group (**Figure 4D-F**).

**Figure 4.**
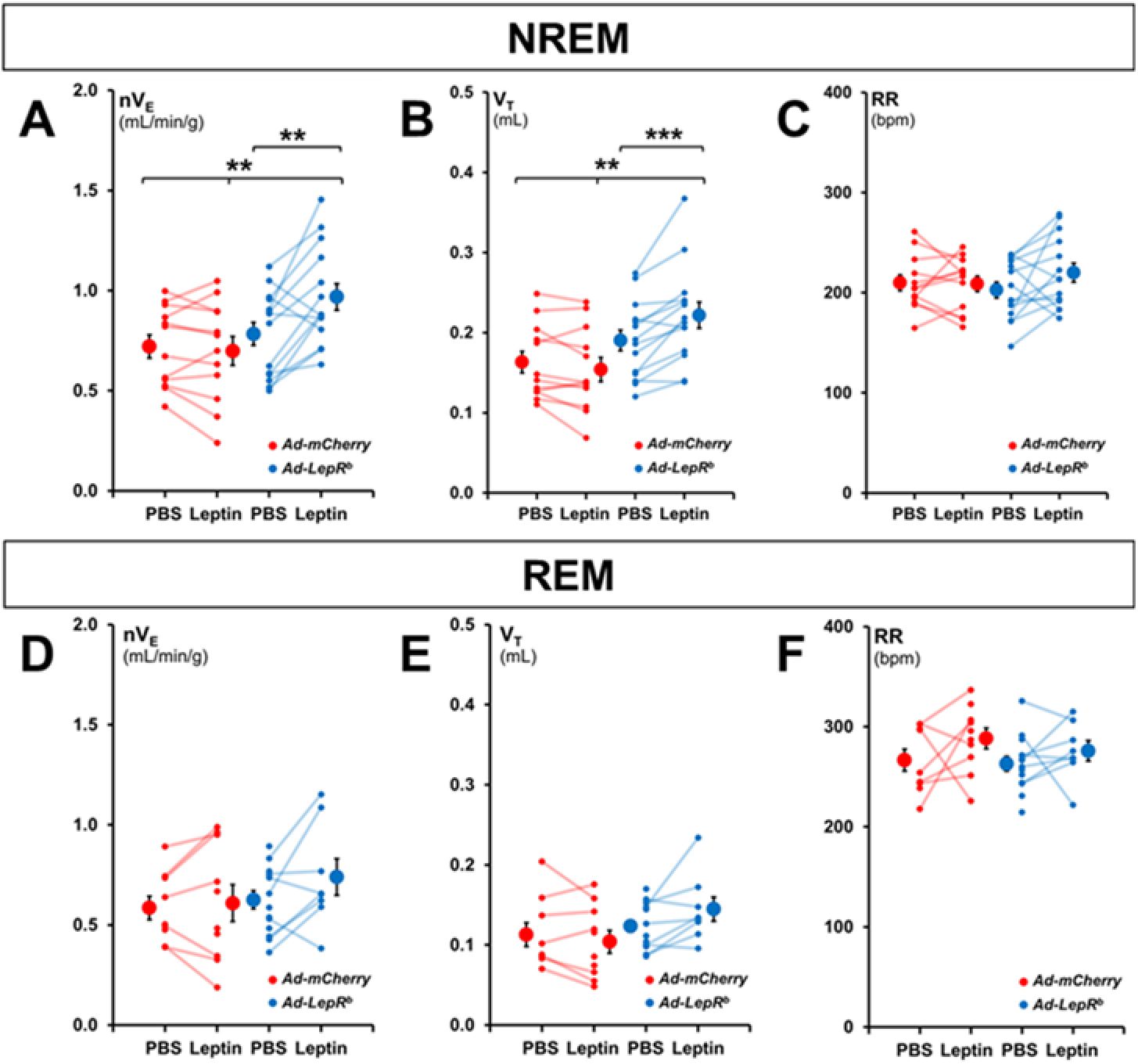
Normalized minute ventilation (nV_E_), tidal volume (V_T_), and respiratory rate (RR) in non-flow limited breathing during (A-C) NREM and (D-F) REM sleep. Each line represents individual mice; mean values ± standard errors are shown. Leptin infusion augmented nV_E_ in Ad-LepR^b^ in NREM sleep compared to PBS injection and Ad-mCherry group, but not in REM sleep. Changes in nV_E_ were caused by an increase in V_T_. **p<0.01; ***p<0.001. NOTE: n = 3 mice transfected with Ad-mCherry and n = 3 transfected with Ad-LepR^b^ did not have REM sleep after PBS injection.

In *Ad-LepR^b^*-transfected mice, ICV leptin increased maximal inspiratory flow (V_I_max) during nonflow limited breathing by 20%, from 3.40 ± 0.25 mL/s to 4.09 ± 0.28 mL/s (**Figure 5A**) and increased mean inspiratory flow rate (MIFR) by 24%,from 1.91 ± 0.14 mL/s to 2.37 ± 0.15 mL/s (**Figure 5B**). Leptin had no significant effect in mice transfected with *Ad-mCherry*. Leptin did not affect V_I_max and MIFR of non-flow limited breaths during REM sleep, regardless of the virus (**Figure 5C,D**).

**Figure 5.**
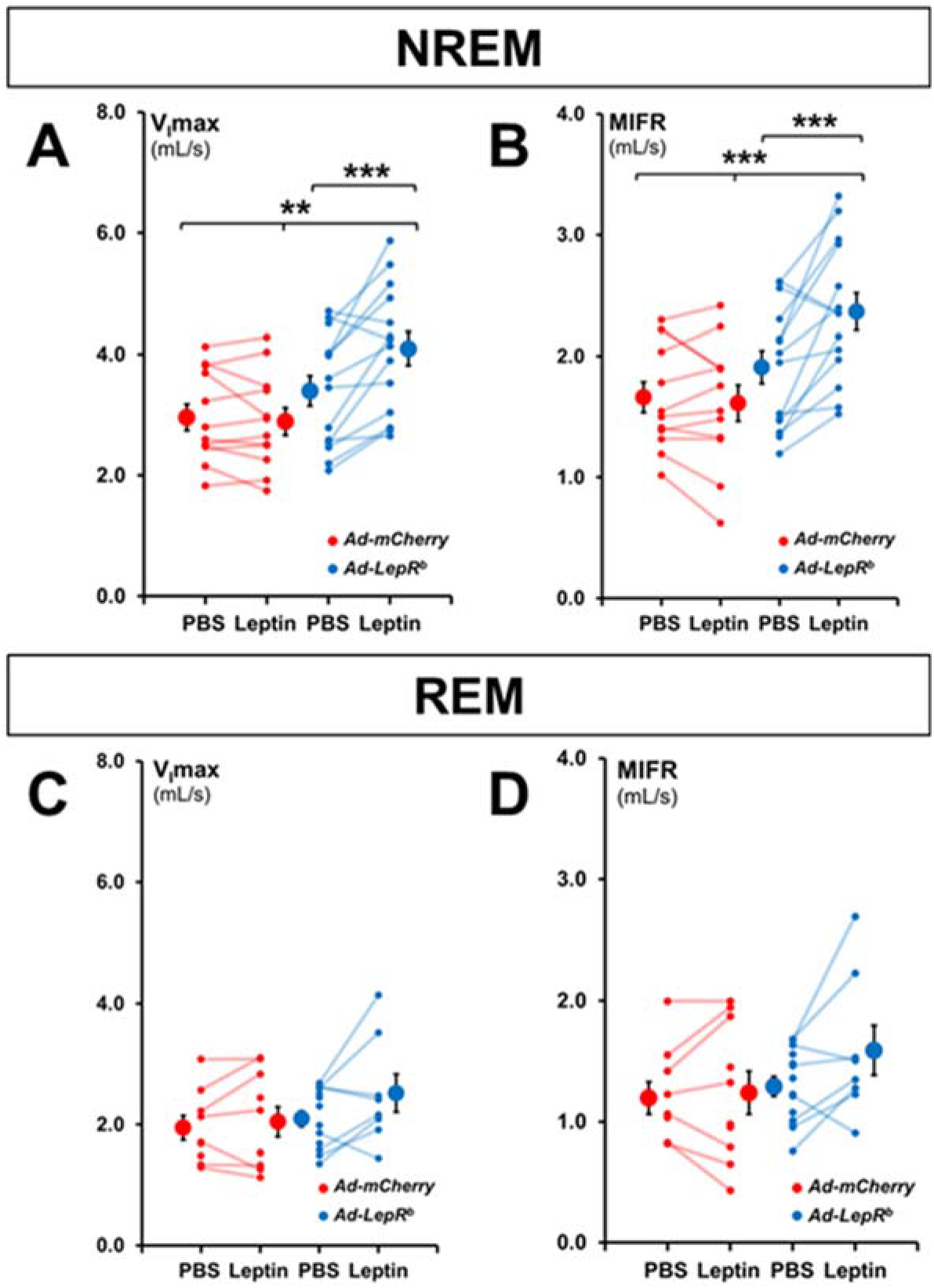
Maximum inspiratory flow (V_I_max) and mean inspiratory flow rate (MIFR) in (A-B) NREM and (C-D) REM sleep during non-flow limited breathing. Each line represents individual mice; mean values ± standard errors are shown. Leptin infusion increased V_I_max and MIFR in NREM in Ad-LepR^b^ mice compared to PBS injection and Ad-mCherry group, but not in REM sleep. **p<0.01; ***p<0.001.

### Effect of leptin on upper airway obstruction in DMH-Ad-LepR^b^ transfected db/db mice

*Db/db* mice developed upper airway obstruction exclusively during REM sleep. The percentage of inspiratory flow limited breaths was negligible for all time points during NREM sleep in both the *Ad-mCherry* and the *Ad-LepR^b^* group. In contrast, upper airway obstruction was observed during REM sleep (**Supplemental Figure 2**). The prevalence of inspiratory flow limited breaths in REM sleep after PBS treatment was the same in the *Ad-mCherry* and the *Ad-LepR^b^* groups. Leptin had no effect on the percentage of inspiratory flow limited breaths in REM sleep in the *Ad-LepR^b^-treated* mice, 14.9 ± 3.4 % of all breaths after PBS and 7.8 ± 2.1 % after leptin. Furthermore, the severity of the upper airway obstruction was not affected either. In the *Ad-LepR^b^* group, V_I_max during obstructed breathing was 1.87 ± 0.12 mL/s after PBS and 2.15 ± 0.22 mL/s after leptin treatment and minute ventilation was 0.62 ± 0.04 mL/min/g after PBS and 0.69 ± 0.08 mL/min/g after leptin, respectively (**Figure 6**).

**Figure 6.**
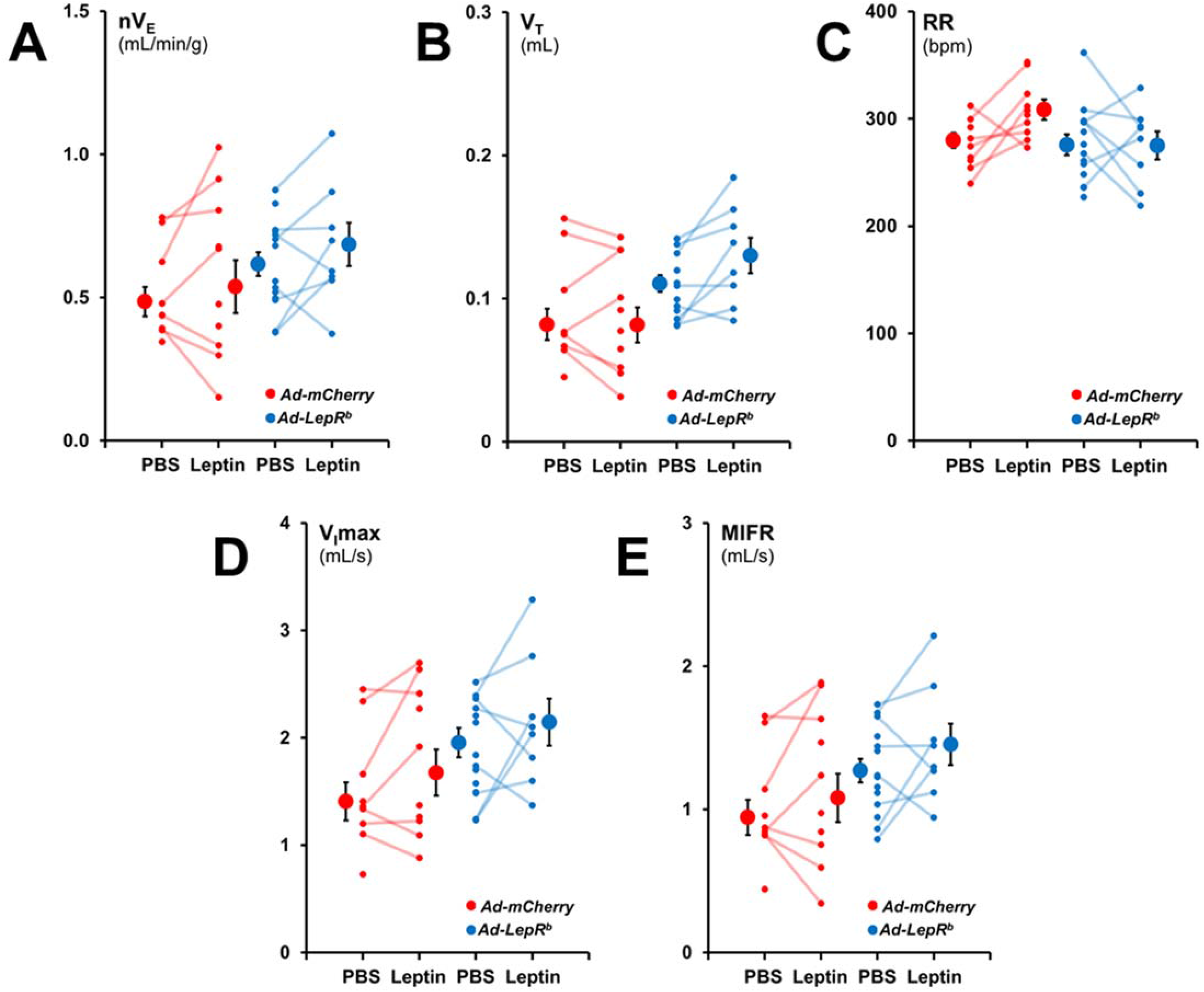
Ventilatory parameters during flow limited breathing in REM sleep. Normalized minute ventilation (nV_E_), tidal volume (V_T_), and respiratory rate (RR), maximum inspiratory flow (VImax), and mean inspiratory flow rate (MIFR). Each line represents individual mice; mean values ± standard errors are shown. No significant effect of LepR^b^ or leptin on any parameters of flow limited breathing was observed.

### Leptin had no effect on metabolism in DMH Ad-LepR^b^ transfected db/db mice

In order to examine if the increase in minute ventilation was due to an increase in CO_2_ production by leptin, we monitored the mice in metabolic cages (**Figure 7**). In mice transfected with *Ad-LepR^b^* in the DMH, ICV leptin did not affect energy expenditure or motor activity. Similar data were obtained in mice transfected with *Ad-mCherry*, except for a curious and inexplicable decrease in VO_2_ with leptin administration.

**Figure 7.**
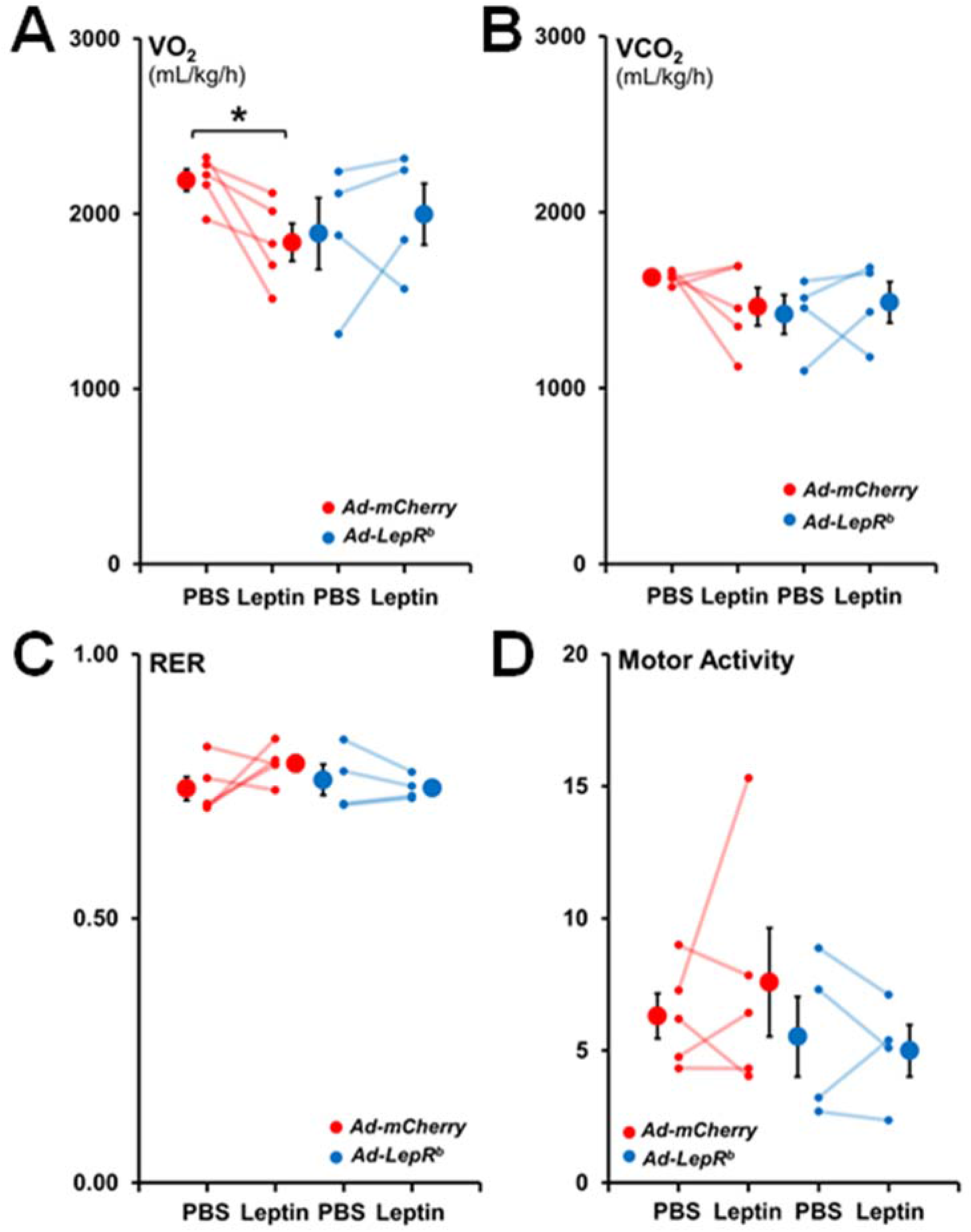
Energy expenditure and motor activity during the light phase in db/db mice infected with Ad-LepR^b^-GFP or Ad-mCherry. Each line represents individual mice; mean values ± standard errors are shown. There were no changes in (A) total oxygen consumption (VO_2_), (B) total carbon dioxide production (VCO_2_), (C) respiratory exchange ratio (RER) and (D) total motor activity between Ad-LepR^b^ and Ad-mCherry with either PBS or leptin infusions. *p<0.05.

## DISCUSSION

The main finding of our study is that targeted expression of LepR^b^ in the DMH in an animal model with SDB (LepR^b-^deficient *db/db* mice) resulted in a beneficial sustained increase in minute ventilation during NREM sleep in response to ICV leptin infusion. We also report several other novel findings. <u>*First*</u>, LepR^b^-deficient obese *db/db* mice exhibit chronic alveolar hypoventilation, which is similar to humans with OHS. <u>*Second*</u>, adenoviral vector-driven DMH transfection of *LepR^b^* resulted in LepR^b^ expression exclusively in neurons, and these neurons were MC4 positive in a pattern similar to *LepR^b^-Cre-GFP* mice. <u>*Third*</u>, leptin-induced hyperventilation mediated by LepR^b^ (+) DMH neurons was not associated with an increase in VCO_2_ implicating stimulation of respiratory control centers. <u>*Fourth*</u>, ICV leptin suppressed REM sleep acting upon LepR^b^ (+) DMH neurons. <u>*Fifth*</u>, *db/db* mice exhibited upper airway obstruction during REM sleep, which was not affected by expression of LepR^b^ in DMH.

### *Db/db* mice exhibit alveolar hypoventilation

Compared to our historic data in lean and diet-induced obese C57BL/6J mice^14^, *db/db* mice had excellent sleep efficiency and had less fragmented NREM sleep in the barometric plethysmography chamber. Total REM sleep time at baseline was similar to diet-obese mice and decreased compared to lean mice^14^ due to a decreased number of REM sleep bouts. Our data provided first demonstration that leptin-resistant obese *db/db* mice hypoventilate chronically, similar to leptin deficient *ob/ob* mice^27^ and leptin-resistant diet-induced obese mice^14^. Awake unrestrained unanesthetized *db/db* mice had compensated respiratory acidosis with PaCO_2_ of 42 mmHg, which was similar to that in *ob/ob* mice^27^ and slightly higher than PaCO_2_ of 39 mmHg previously reported in diet-induced obese mice^14^. In contrast, lean mice on the same genetic background had PaCO_2_ of 30-33 mmHg^14,27^. Sleep studies in *db/db* mice demonstrated hypoventilation during NREM sleep with average nV_E_ of 0.78 ml/min/g, compared to our historic data of 1.2-1.3 mL/min/g in lean mice^14^, as a cause of CO_2_ retention. *ob/ob* and diet-induced obese mice showed the same levels of hypoventilation as *db/db* mice^14,18,19^. However, *ob/ob* and *db/db* mice breathe at lower V_T_ and higher RR than diet-induced obese mice (~ 0.19 *vs* 0.23 mL and > 200 *vs* 150 breaths per minute respectively), which could lead to higher dead space ventilation and lower alveolar ventilation resulting in higher CO_2_ levels observed in our studies^14,18,19,27^. Thus, leptin-resistant *db/db* and diet-induced obese mice as well as leptin-deficient *ob/ob* mice exhibit obesity hypoventilation, similar to obese humans with OHS^8,50^. OHS leads to high mortality with 23% untreated patients dying during the 18-month observation period^8,51^. Understanding of mechanisms by which leptin acts on respiratory centers in the brain to increase ventilation is important for future therapy of SDB.

### The effect of leptin on control of breathing and the role of LepR^b^ signaling in DMH MC4 (+) neurons

Leptin-LepR^b^ signaling in the DMH increases metabolic rate by activating thermogenesis in brown adipose tissue^52,53^. In our study, administration of leptin ICV to LepR^b^-deficient *db/db* mice expressing LepR^b^ receptor exclusively in DMH induced a modest increase in core body temperature of 0.5°C (**Table 1**), but there was no significant changes in VCO_2_ or VO_2_ (**Figure 6**). The lack of metabolic response could be attributed to relatively low levels of LepR^b^ expression, which was detected only in 11.6% of DMH cells. Nevertheless, even this low level of LepR^b^ expression restored sensitivity to respiratory effects of leptin. ICV leptin increased minute ventilation in NREM sleep without concurrent increases in VCO_2_ or physical activity, which suggests stimulation of respiratory control centers in the brain. In fact, the magnitude of the effect of leptin signaling in DMH on minute ventilation in our current study was identical to increases induced by leptin *via* signaling in the entire brain of *ob/ob* mice and diet-induced obese mice in our previous studies^19,20,54^.

Bassi *et al* microinjected leptin into the ventrolateral medulla of *ob/ob* mice, specifically into the retrotrapezoid nucleus/parafacial respiratory group (RTN/pFRG)^55^, a primary site for central chemoreception^56–58^, and found an increase in the hypercapnic ventilatory response (HCVR) in awake animals. Another group found that microinjection of leptin into the nucleus of the solitary tract (NTS) significantly increased respiratory activity in anesthetized rats under hypercapnic conditions^59^. LepR^b^ (+) DMH neurons may stimulate breathing *via* their projections to these medullary nuclei, but overall mechanisms of respiratory effects of leptin in DMH remain unknown.

Another intriguing finding of our study was that *Ad-LepR^b^* transfection resulted in LepR^b^ expression in MC4 (+) neurons of DMH (**Figure 2**). Furthermore, MC4 staining of DMH in *LepR^b^-Cre-GFP* mice showed a similar pattern. MC4 activity is regulated by leptin^60,61^. MC4 signaling increases metabolic rate by inducing metabolic uncoupling in brown adipose tissue^62^. The melanocortin system is also involved in respiratory effects of leptin. Mice producing agouti-related peptide with deficient leptin and melanocortin signaling have suppressed hypercapnic ventilatory response^63^. An MC4 blocker SHU abolished leptin-induced respiratory chemosensitivity^64^. Thus, MC4 signaling may play a role in mediating respiratory effects of leptin signaling in DMH.

### The effect of leptin on upper airway patency during sleep

Our data also showed that *db/db* mice exhibit mild inspiratory flow limitation at baseline, exclusively in REM sleep. We have previously reported very low prevalence of obstructed breaths in NREM sleep in *ob/ob* mice (<1%-3%), whereas prevalence of obstructed breaths in REM sleep was higher^18,19^. More severe impairment of upper airway patency could be related to older age or higher body weight of *ob/ob* mice in the previous studies, 65-70 g^18,19^ compared to 47-50 g in *db/db* mice in the current study (**Table 1**). In contrast, diet-induced obese mice showed much higher prevalence of obstructed breathing in both NREM and REM sleep, 13-15% and 35-45% respectively, despite having similar weight to *db/db* mice^14,20^. Thus, diet-induced obesity appears to predispose to OSA compared to genetic models of leptin deficiency and resistance.

Our previous work demonstrated that leptin acts in the brain to relieve upper airway obstruction during sleep, both in leptin-deficient *ob/ob* mice^19^ and in leptin-resistant diet-induced obese animals^20^. LepR^b^ was not detected in the hypoglossal motoneurons, but there was evidence that LepR^b^ positive neurons are synaptically connected to hypoglossal neurons^20^. Moreover, leptin activated hypoglossal motoneurons in brain slices^54^. Indirect evidence indicated that leptin acts in the forebrain, possibly in DMH, to regulate upper airway patency during sleep^19^. Our current study did not confirm this report showing that leptin had no effect on flow limited breathing in *db/db* mice expressing LepR^b^ in DMH neurons. However, leptin substantially shortened REM sleep, from 5.2 to 1 min and the prevalence of upper airway obstruction at control conditions was only 14.9%. Thus, our data on the role of leptin signaling in DMH in the pathogenesis of OSA are inconclusive.

### Leptin and sleep

We have previously reported that ICV leptin shortened REM sleep in *ob/ob* mice^19^, which was consistent with previous data in rats^65^. In humans, a sharp decline in leptin levels through the night was associated with REM sleep rebound^66^. Taken together, this data suggests that leptin shortens REM sleep, possibly *via* LepR^b^ signaling in the DMH. The role of DMH in REM sleep regulation is insufficiently studied. Several hypothalamic mediators, including melanin concentrating hormone^67^ and hypocretin^68–70^, may impact REM sleep, but neurons producing these molecules were located predominantly to the lateral hypothalamus. Chen *et al* showed that galanin-expressing GABAergic neurons in the DMH constitute two different neuronal populations with opposing effects on REM sleep^71^, but the relevance of this finding for our study is uncertain. Finally, LepR^b^ signaling in DMH may decrease REM sleep indirectly by increasing body temperature^72^, but the effect of leptin on body temperature in our study was modest.

### Limitations of the study

Our study had a number of limitations. <u>*First*</u>, mice were heavily instrumented and sleep was recorded only for six hours after ICV injections, which could affect the quality of sleep. In order to account for these inevitable confounders we compared PBS with leptin within the same mice and *Ad-LepR^b^* with control virus. <u>*Second*</u>, an adenoviral vector may induce non-specific adverse effects in mice. In order to counter this problem we used control *Ad-Cherry* vector and completed all studies within 13-20 days after transfection. Furthermore, we found that *Ad-LepR^b^* by itself did not induce any metabolic or respiratory effects prior to leptin injections. *Third, db/db* mice have very high plasma leptin levels at baseline^14^, nevertheless ICV delivery of supplemental leptin was required to induce respiratory effects. <u>*Fourth*</u>, although our model allowed us to identify effects of leptin on control of breathing, *db/db* mice appeared to be a suboptimal model to study effects of leptin on upper airway during sleep. Given that mice with diet-induced obesity showed more severe upper airway obstruction during both NREM and REM sleep^14,20^ than *ob/ob*^18,19^ and *db/db* mice, chemogenetic and/or optogenetic approaches in LepR^b^-Cre diet-induced obese mice may be more promising for OSA research. <u>*Fifth*</u>, the goal of our study was to establish if LepR^b^ DMH neurons are involved in control of breathing, but we did not examine synaptic projections of these neurons to downstream respiratory control centers in the brainstem.

## CONCLUSION

Leptin stimulates ventilation by acting on LepR^b^ (+) neurons in DMH and this effect is independent of metabolism. Further elucidation of molecular make up of LepR^b^ (+) DMH neurons and their projections to respiratory control centers may lead to new therapies for OHS, a common and lethal condition, and novel treatment strategies for other forms of hypoventilation characterized by diminished CNS output.

## ACKNOWLEDGEMENTS

This study was supported by the grants from the National Institutes of Health R01 HL128970, R01 HL133100, R01 HL13892, and R61 HL156240 (all to VYP), and American Heart Association Career Development Awards 19CDA34700025 (MKS) and 19CDA34660245 (TFC).

## SUPPLEMENTAL

**Supplemental Figure 1.**
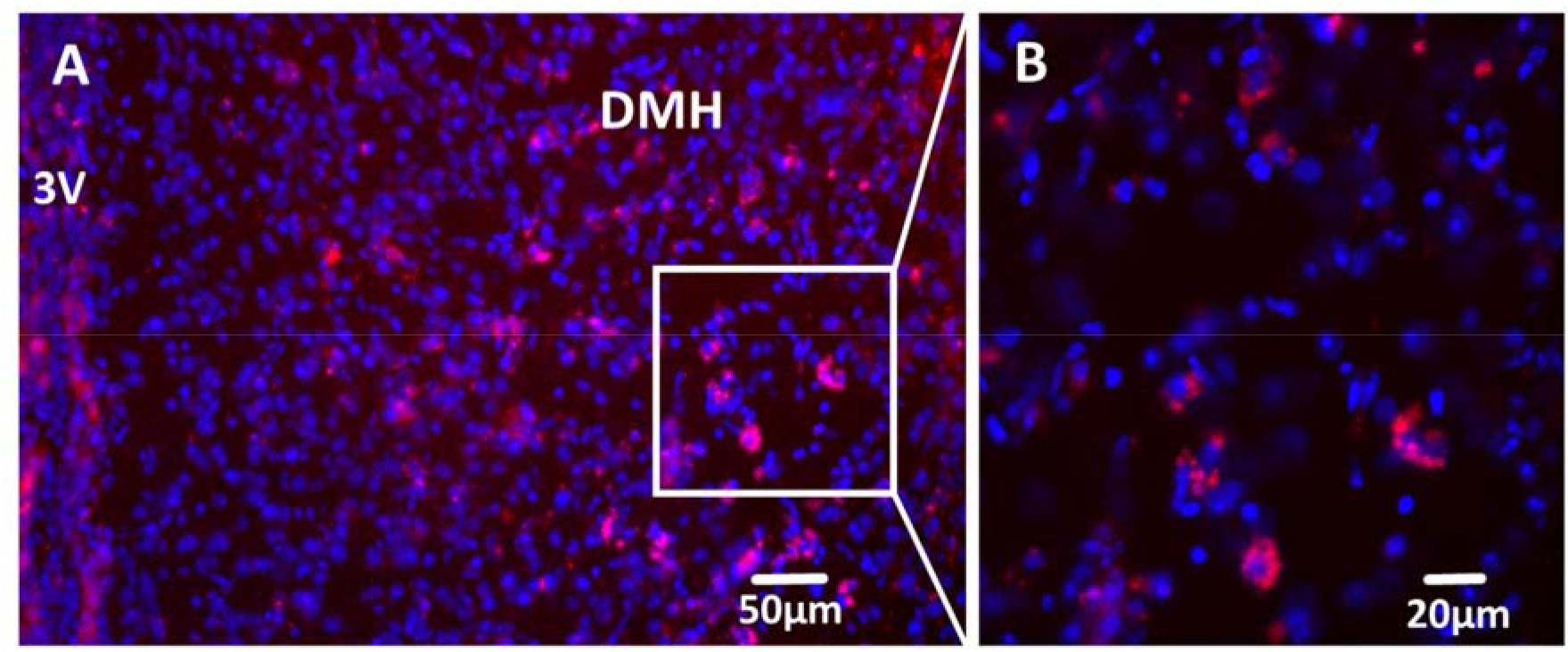
Localization of mCherry in DMH of db/db mice after Ad-mCherry stereotactic injection. (A) mCherry (in RED) was expressed in DMH of db/db mice 9 days after Ad-mCherry infection; staining of nuclei with DAPI (4’,6-diamidino-2-phenylindole) in BLUE. (B) same at higher magnification.

**Supplemental Figure 2.**
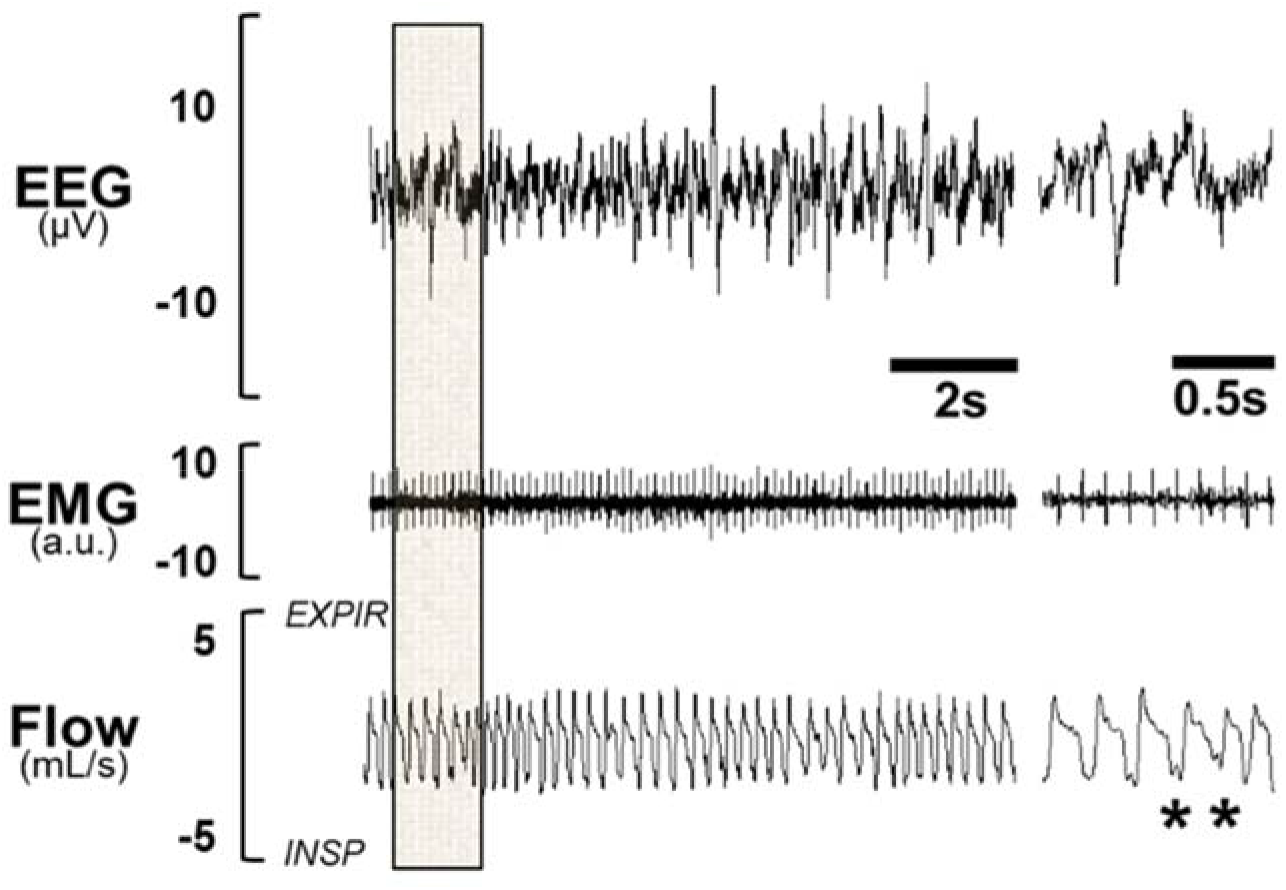
Representative REM sleep recording in db/db mice. Inspiratory flow limitation is present in breaths (indicated by *). REM sleep in Ad-mCherry transfected group showed no significant difference when compared to the Ad-LepR^b^ group. Leptin had no effect on the percentage of obstructed breaths in REM sleep in the Ad-LepR^b^-treated mice, 14.9 ± 3.4 % of all breaths after PBS and 7.8 ± 2.1 % after leptin.

